# Revisiting Feature Selection with Data Complexity

**DOI:** 10.1101/754630

**Authors:** Ngan Thi Dong, Megha Khosla

**Affiliations:** L3S Research Center, Leibniz University Hannover, Germany

## Abstract

The identification of biomarkers or predictive features that are indicative of a specific biological or disease state is a major research topic in biomedical applications. Several feature selection(FS) methods ranging from simple univariate methods to recent deep-learning methods have been proposed to select a minimal set of the most predictive features. However, there still lacks the answer to the question of “which method to use when”. In this paper, we study the performance of feature selection methods with respect to the underlying datasets’ statistics and their data complexity measures. We perform a comparative study of 11 feature selection methods over 27 publicly available datasets evaluated over a range of number of selected features using classification as the downstream task. We take the first step towards understanding the FS method’s performance from the viewpoint of data complexity. Specifically, we (empirically) show that as regard to classification, the performance of all studied feature selection methods is highly correlated with the error rate of a nearest neighbor based classifier. We also argue about the non-suitability of studied complexity measures to determine the optimal number of relevant features. While looking closely at several other aspects, we also provide recommendations for choosing a particular FS method for a given dataset.

## 1 Introduction

One of the core issues in applying machine learning and data mining techniques to biomedical domain is the so called *curse of dimensionality*. This refers to the phenomena largely observed in biomedical data: small number of instances with high dimensionality (features), leading to high sparsity in data, which adversely affects algorithms designed for low-dimensional space. In addition, with a large number of features, learning models tend to overfit hence leading to a drop in performance on unseen data. Consider for example, gene micro-array analysis, where data might contain thousands of variables in which many of them could be exceedingly correlated. Generally, for a pair of perfectly correlated features, keeping one is sufficient to retain the descriptive power of the pair. These redundant but relevant features can contribute significantly to the over-fitting of a model. In addition, there could exist some noisy features (e.g, the ones having no correlation to the class) leading to erroneous class separation. In such cases, feature selection, as a dimensionality reduction technique, has proven to be effective in preparing the data or selecting the most relevant features for performing downstream machine learning tasks such as classification. In addition, it plays a critical role in *biomarker discovery* for diagnosis and treatment of complex diseases.

Feature selection (FS) has been widely applied in bioinformatics [11, 38, 39, 22] and can be broadly classified into filter, wrapper and embedded methods. While filter methods evaluate the relevance of features by considering only the intrinsic properties of the data, the wrapper method selects a feature subset by iteratively selecting features based on the classifier performance. The embedded methods combines feature selection and classifier construction using an integrated model building process. Feature selection has attracted strong research interest in the past several decades and a huge number of methods have been proposed. Nevertheless, the main question of *which FS method to use when* remains unanswered. In this work we study the above problem from the perspective of data complexity and instead ask *if data complexity measures[4] can be leveraged to prefer a particular method*.

We conduct a comprehensive empirical comparative study using 11 FS methods, including representatives of (i) filter, (ii) wrapper, (iii) embedded (iv) and recently proposed deep learning approaches on 27 biological datasets with varying properties. Table 1 presents a summary of those 11 selected techniques. In particular we relate the performance difference of various methods to the dataset properties determined by several data complexity [39, 4, 25] measures as explained in Section 2. To the best of our knowledge, this is the first work which applies data complexity measures to understand the suitability of FS methods. Our results show that the difficulty of finding the most relevant features for all methods is correlated with a very easy to compute data complexity measure which corresponds to the estimated error rate of a nearest neighbor based classifier. While this might seem very unsurprising, we also find methods which are more affected by it than others. Intuitively, the high correlation with such an error rate also implies that the FS method was not able to extract relevant features using which samples sharing the same class could be put closer.

**Table 1:** List of evaluated FS methods with link to used implementation.

| Method Name | Type | Link |
| --- | --- | --- |
| INFO GAIN | filter, supervised | link |
| CHI-SQUARE | filter, supervised | link |
| RELIEFF | filter, supervised | link |
| MULTI-CLUSTER FEATURE SELECTION (MCFS) | filter, unsupervised | link |
| LASSO | embedded, supervised | link |
| ELASTIC NET | embedded, supervised | link |
| HSIC-LASSO | embedded, supervised | link |
| BORUTA | wrapper, supervised | link |
| MULTILAYER PERCEPTRONS (MLP) | deep learning, supervised | link |
| STACKED CONTRACTIVE AUTOENCODER (SCA) | deep learning, supervised | link |
| DEEP BELIEF NETWORK (DBN) | deep learning, supervised | link |

One of the issues in evaluating FS methods is that, in most of the cases, the optimal size of important or non-redundant features is not known. In particular, an FS method returns either a subset of features or a list of ranked/weighted features. For ranking methods, the higher the weight/rank, the more important the feature is. For methods which return a subset of features, all features in the returned list are considered important. Most of the previous works merely cover a small range of pre-specified selected features when evaluating the suitability of a method. Yet it could also happen that a given method reaches its peak performance with a significantly smaller number of features. For example, peak performance might be reached with 1% of the top ranked/weighted features. Adding additional features could degrade its performance considerably. This reveals an issue in ranking, where noisy features are erroneously ranked higher. In this work, we experimented using a wider range of selected features. We found out that the optimal number of features vary for different methods and cannot be predicted by using any of the presented data complexity measures. Moreover there is no monotonic trend observed with increasing number of selected features and performance which calls into question the goodness of an FS method whose performance might have only be evaluated using only a specific number of features. In particular, we also did not find a correlation between the optimal number of features predicted by performing PCA (while preserving a high percentage of data variability) and the actual number of features for which a particular method obtains its maximum performance (on classification task).

Summarizing our findings, we provide a priority list to choose one method over the other based on the dataset characteristics and properties of the method.

### 1.1 Related Work

In this section we provide a brief overview of related reviews and comparative studies and their differences to the present work. We also point to various works which have studied data complexity measures either to quantify difficulty in classification or deciding cut-off thresholds for feature selection methods.

Degenhardt et al. [11] studied and compared various Random Forest based methods on two high dimensional real word biological datasets with respect to classification performance, stability and run time. But their focus is limited to a particular type of methods. Moreover, the number of datasets considered is also quite small to be able to generalize the results.

Taking a broader perspective, Neto et al. [26] construct a large scale study on simulated data to investigate the effects of sample size, number of features, true model sparsity, signal-to-noise ratio, and feature correlation on predictive performance of ridge regression, Elastic Net and Lasso methods. Through diverse, carefully designed experiments, they focus on the strengths and weaknesses of only those three methods under very particular conditions.

Urbanowicz et al. [38] evaluated 13 existing and their proposed 3 Relief-Based algorithms for a genetic simulation study. They run experiments on 2280 simulated datasets cover a wide range of problems and types. Nevertheless, the question of whether the findings can be applied to the real-world datasets is still left open.

Wang and Barbu [39] try to answer the questions of (i) whether filter methods help improve classification model and (ii) how existing filter methods are different from each other in terms of predictive capabilities. They construct experiments on five regression and five classification datasets. They measure the classification performance of FS methods on 40 different runs with 30 different numbers of selected features. In addition to limitation to filter methods, the focus is also not primarily on biological datasets. The survey conducted by Li et al. [22] seems to be the most comprehensive one, which presented a general summary of existing works on FS methods from different data type perspectives. Nevertheless, deep learning methods are not included in their study.

Data complexity measures for feature overlap are used in [30] to choose the feature cut-off threshold for ensemble FS methods. We argue that their problem statement is different from ours as (1) their work is focus solely on ensemble FS methods, (2) the complexity measures were used to guide the aggregation of multiple feature subsets returned by multiple FS methods, not the number of selected features for individual method and (3) they only experimented with 6 DNA binary microarray datasets which is too small to derive any conclusion. In [18], complexity measures were used to quantify the difficulty of classification on two different gene expression datasets. Another work [10] relates complexity measurements to the classification performance of Support Vector Machines on cancer gene expression data.

We conclude that previous works are either (i) too narrow, focusing on a particular class of FS methods or/and using only simulated datasets, or are (ii) too broad, meaning that their comparisons are not focused solely on biological data. In addition, none of these works compare deep learning methods. From the data complexity perspective, none of the works study our proposed problem, i.e., whether one can use a set of complexity measures to guide the choice of a particular feature selection method.

In the next sections, we present details about the complexity measures used and FS methods respectively in sections 2 and 3. Finally, we present our experimental set-up and results followed by conclusion and a priority list on the choice of feature selection methods.

## 2 Data Complexity Measures

In this section, we briefly describe the data complexity measures that we use in our analysis. Data complexity measures [4] have been traditionally used to study the intrinsic difficulty of a classification task on a given dataset. In this work, we relate the data complexity characterized by 4 such measures to the suitability of a particular FS method. From now on, we use *m* to denote the number of features, *n* to denote the number of samples and *n*_*c*_ to denotes the number of classes. The definitions of these measures have been adapted from [25]. We use the ECoL package^2^ to calculate the data complexity measures for the listed datasets.

1. **Error Rate of Nearest Neighbor (NN) classifier (N3):** N3 is measured by the error rate of the 1-NN classifier using leave-one-out cross validation. Formally, 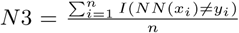, where *NN* (*x*_*i*_) is the predicted target value for sample *x*_*i*_ using all other samples as the training set. High N3 scores indicate that instances of different classes are close together.
2. **Ratio of the PCA dimension to the original dimension (T4) [24]**: T4 is the ratio of the number of PCA (Principal Component Analysis) components needed to represent 95% of data variability on the total number of features. Higher T4 scores indicate a larger portion of the original features set is necessary to preserve the information of the dataset, thus, implying the need to use a larger number of features for a given task.
3. **Sparsity (T2):** Sparsity is defined as 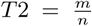. Highly sparse datasets can be difficult for classification since learning process can be hindered in the low density regions.
4. **Class Imbalance(C2):** The imbalance ratio measures the differences in the number of instances per class in the dataset and is computed as:

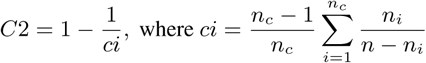

where *n*_*i*_ is the number of samples in class *i* and *n*_*c*_ is the number of classes. Higher values of C2 indicate higher class imbalance.

## 3 Compared Feature Selection Methods

In this work, we compare representatives of a wide range of unsupervised and supervised FS methods, including filter, embedded, wrapper and deep learning (DL) based methods. The choice of our models is based on recommendations from previous works, as well as our own initial set of experiments that we conducted. We included a larger number of methods from each category, from which we choose a subset of the best-performing approaches. Unlike previous evaluation works, we have included recent deep learning methods in our study. A summary of the compared methods is provided in Table 1.

### Info gain

Information gain (Info gain)[29] measures the amount of information in bits about the class prediction, assuming that the only information available is the presence of a feature and the corresponding class distribution. The information gain from splitting the data set(*S*) using the values of the feature *f*_*i*_ is given by

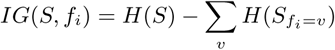

where the entropy *H*(*S*) = Σ_*C*_ *p*(*S, C*) log *p*(*S, C*). *p*(*S, C*) denotes the probability that a training example in *S* belongs to class *C*. The notation 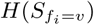 corresponds to the entropy of the dataset after fixing the value of feature *f*_*i*_ to *v*. The features can be ranked based on the information gain scores, higher the information gain, the more important a feature is.

### Chi-square [41]

Chi-square feature selection method utilizes the test of independence to assess whether the feature is independent of the class label. It iteratively calculates the chi-square statistics between each feature with the target class label. If these two variables (feature and target variables) are independent then we eliminate that feature from the feature set since it contributes nothing to the prediction of the target variable. The smaller the *p*-value (corresponding to chi-square test), the more is important the feature.

### ReliefF [20]

ReliefF is based on the Relief algorithm whose main idea is to estimate features’ importance according to how well their values distinguish between neighboring samples. Each feature is weighted according to the relationship of *n* random samples to their nearest neighbor(s). For a given sample, RELIEFF selects *k* nearest samples (hits) from the same class and *k* nearest samples (misses) from each of the other classes.

#### Multi-Cluster Feature Selection (MCFS) [6]

MCFS aims to select the most informative features by selecting features, which preserve the clustering structure of the data. It works in two stages. The first stage is responsible for constructing a *k*-nearest neighbor weighted graph from the dataset as well as learning a low-dimensional representation (embedding) for each node (instance) by solving the generalized eigen problem. The second stage is subjected to extracting the importance of each feature by solving a **L1 regularized least squares problem**, such that the clustering structure of the data is preserved.

#### Least Absolute Shrinkage and Selection Operator (Lasso)[36]

LASSO allows feature selection based on the assumption of linear dependency between input features and output values and use **L1-penalty** (regularization) in the final loss function. With respect to classification, this translates to presence of linear decision surface separating the two classes. For datasets with binary classes, with input training data *X* ∈ ℝ^*n×m*^ and target (class) variable *y* ∈ 1{-1, 1}^*n*^, we seek **w** ∈ ℝ^*m*^ that minimizes *L*1-regularized objective function:

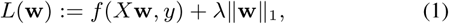

where 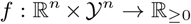 is a loss function, and *λ* ∈ ℝ_*≥*0_ is a regularization parameter,*w*_*j*_ is the weight coefficient corresponding to feature *j*. Features with non-zero weight coefficients are considered important features by Lasso. However, for a set of correlated features, Lasso tends to randomly pick only one feature. In this work, we use Lasso for feature selection in datasets with two classes and use logistic loss for classification loss function *f*.

### Elastic Net [43]

Elastic Net addresses the drawback of Lasso, by incorporating both the L1 and L2 regularization penalties. Like Lasso, Elastic Net simultaneously produces a model and performs automatic variable selection via shrinkage; however, it is also able to account for subsets of correlated features by using additionally the **L2 regularization** term (*λ*_2_||**w**||_2_) in its optimization function. We apply Elastic NET for feature selection in datasets with 2 classes with logistic regression for determining the classification loss.

### HSIC-Lasso [40]

Hsic-Lasso extends Lasso by finding nonlinear feature dependencies. In particular, it finds non-redundant features with strong statistical dependence on the output classes using kernel-based independence measures such as the Hilbert-Schmidt independence criterion (HSIC). The optimization function for Hsic-Lasso is obtained from (1) by using particular forms of universal reproducing kernels[40] for feature and target variable transformations. Like Lasso it also employs L1-regularization.

### BORUTA[21]

The key idea behind this approach is to compare the importance of every feature with those of random or *shadow* variables using statistical testing and several runs of Random Forest. A shadow variable is created for every feature by permuting its original value. After that, a Random Forest classifier model is trained on the extended dataset, while the importance scores/weights of all of the attributes, including the shadow variables, are calculated at the same time. Since the shadow variables are designed to be random, their weights are expected to be close to zero. Boruta uses the highest importance score of all shadow variables as a threshold to determine whether a feature is truly important or redundant.

#### Deep Feature Selection (DFS) Model [23]

Li et al.[23] constructs a DFS model from an multilayer perceptron **(MLP)** by adding a sparse one-to-one linear layer between its input layer and the first hidden layer. The weights of this one-to one layer are considered as the importance of the corresponding features. The model parameters (including those of the one-to-one layer) are trained using the negative log likelihood loss function (cross entropy for multi-class classification), along with an **Elastic Net based regularization** for the feature importance weights as well as other parameters of the network. In addition, the authors also experiment by replacing MLP with stacked contractive autoencoders(SCA) and Deep Belief Networks(DBN) which we also include in our experiments.

1. **Stacked contractive autoencoder (SCA)** SCA fundamental building block is a stack of contractive auto-encoders. A contractive auto-encoder [28] is a type of auto-encoder with the addition of Frobenious norm over the parameters in its loss function. The Frobenious norm is believed to help make the model more robust to small changes in the input. As in [23] we experiment by replacing MLP with SCA in the DFS model.
2. **Deep belief network (DBN) [17]** DBN basically employs the same architecture as MLP but instead of densely connected hidden layers, DBN uses a stack of Restricted Boltzmann Machines (RBMs). Again, as in [23] we experiment by replacing MLP with DBN in the DFS model.

## 4 Evaluation Set Up

A summary of the compared FS methods is given in Table 1 and the datasets with their statistics are summarized in Table 2. For each dataset, we fill the missing values (if any) with the nearest neighbor values, then we use z-score^3^ to normalize the feature values. After that, we run 6 times five-fold cross validation on the dataset with different random states. We collect results from 30 runs to get a close and trustworthy estimate of each method performance. We choose the number of selected features from 20 to 200 with a step of 20. There are several reasons for investigating over a wider range of selected features. **First**, for different number of selected features, FS methods show a varying performance. We also observed that performance does not always show a monotonic relation with the number of selected features. As we do not know apriori what is the optimal/best number of selected features and different users might choose different number of selected features for their datasets. Thus, we argue that constructing and comparing the results over a range of selected features is more meaningful than just a fixed number of selected feature. **Second**, we want to compare the performance of different methods at different points to see how their performance change with regards to the number of selected features? Can one observe any monotonic trend? **Do the experimented methods reach their peak performance for the same number of optimal features?** Can we infer any thing from those peak performance points?

**Table 2:**
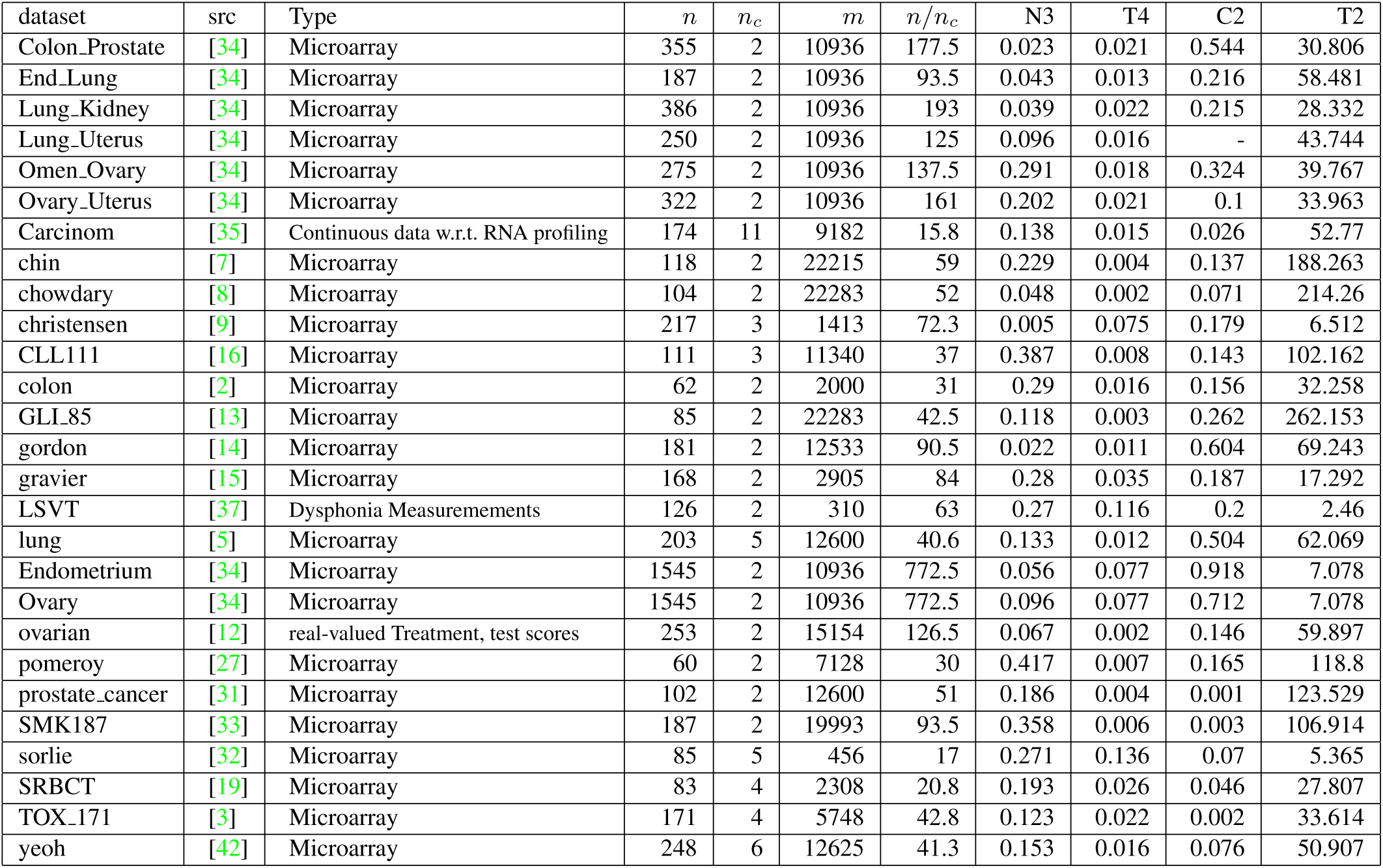
Dataset source, statistics and complexity measures with *n*: number of samples, *n*_*c*_: number of classes, *m*: number of features.

The returned relevant features were then used to train the classification model using the training data. The performance was then tested on the test fold using only the returned relevant features (on training set). For classification, we use a Multilayer Perceptron classifier model^4^. We report the average *F*_1_ scores (harmonic mean of precision and recall) as the measurement for a feature selection method classification performance. We use the same vanilla set up with default parameters for all of the feature selection algorithms and learning models to train the classifier.

We use Pearson correlation coefficient ^5^,^6^ to calculate the correlation between a feature selection method performance and the data complexity scores. We **normalize complexity scores over the datasets with z-score** normalization before measuring correlations. We only report correlation with p-value smaller or equal to 0.05 (statistical significance level of 95%). We point out that comparing the methods based on their space and time complexities is out of scope of this work.

## 5 Results

In Table 2, we present the complexity measurements for all of the datasets. From the table we can see a wide range of selected datasets whose number of samples range from dozens to over more than a thousand, the number of feature range from several hundreds to more than twenty thousands. We also select datasets for both binary and multi-class classification tasks. The range of average number of samples per class (*n/n*_*c*_) range from below twenty to over seven hundreds.

### 5.1 Performance with regards to data complexity

Table 3 present a summary of correlation between the selected FS methods with regards to the presented data statistic and complexity scores. A dash (-) in the table indicates a non-significant correlation (*p* > 0.05). An interesting fact that we discover is that only some of the FS methods performance correlate with the number of average training samples per class (*n/n*_*c*_). Instead, we find those FS methods performance is highly and consistently correlated with N3 - the error rate of the nearest neighbor classifier. It turns out that the smaller the error rate, the better the feature selection method classification performance. In terms of standard deviation, the higher N3, the higher is the variance of FS methods performances over different runs and set up. In the first glance this result might not look surprising. After all, N3 denotes the error rate of a very simple classifier. But we argue that in principle the feature selection methods should have been able to overcome this posed hardness by selecting a subset of features such that closer neighbors have same classes. This seems to be not the case given that the datasets with higher N3 (computed using all features) is still correlated with classification performance over a selected subset of more relevant features. Moreover, the absolute correlation values are different for different methods. We can leverage this information to prefer one method over the other for harder datasets which show a large error rate with a simple 1-NN classifier (when using all the features).

**Table 3:** Statistically Significant Correlation between FS methods performance and the datasets’ characteristics with *p* values *p* <= 0.05. Entries marked *−* have correspond to *p* values > 0.05

| method | others | N3 | C2 |
| --- | --- | --- | --- |
| CHI-SQUARE | - | -0.72 | 0.42 |
| LASSO | - | -0.87 | - |
| INFO GAIN | - | -0.71 | - |
| ELASTIC NET | - | -0.81 | - |
| RELIEFF | - | -0.7 | - |
| BORUTA | - | -0.69 | - |
| DBN | - | -0.67 | - |
| SCA | - | -0.61 | - |
| MLP | - | -0.67 | - |
| MCFS | - | -0.72 | - |
| HSIC-LASSO | - | -0.74 | - |

**Table 4:** Correlation between the standard deviation of FS methods performance and the datasets’ characteristics with *p* <= 0.05

| method | others | $n$ | N3 | T2 | C2 | $n/n_c$ |
| --- | --- | --- | --- | --- | --- | --- |
| CHI-SQUARE | - | -0.49 | 0.57 | - | -0.55 | -0.51 |
| LASSO | - | -0.57 | 0.77 | - | -0.5 | -0.57 |
| INFO GAIN | - | -0.39 | 0.64 | - | -0.43 | -0.39 |
| ELASTIC NET | - | -0.61 | 0.65 | 0.49 | -0.54 | -0.61 |
| RELIEFF | - | - | 0.65 | - | -0.42 | - |
| BORUTA | - | -0.41 | 0.68 | - | -0.46 | -0.41 |
| DBN | - | -0.5 | 0.55 | - | -0.58 | -0.52 |
| SCA | - | -0.44 | 0.62 | - | -0.49 | -0.46 |
| MLP | - | -0.44 | 0.69 | - | -0.45 | -0.45 |
| MCFS | - | -0.5 | 0.64 | - | -0.54 | -0.51 |
| HSIC-LASSO | - | - | 0.57 | - | - | - |

### 5.2 Feature selection methods average performance rank

For each dataset, we calculate the average performance of each method over the range of selected features. Figure 2 present those average F1 scores. There exist blanks because we only run Lasso and Elastic Net for binary classification problems. From those values, for each dataset, we sort them in descending order to get the rank for each FS method. Figure 4 gives details about the rank of each FS method based on average performance over the range of selected datasets. Figure 1 presents the average performance rank for each feature selection methods over all datasets. Looking at the plots we can see that on average Info gain ranks the highest and is also the method with lowest variance in performance. Hsic-Lasso comes second in terms of both for performance and variance followed by ReliefF and MLP.

**Figure 1:**
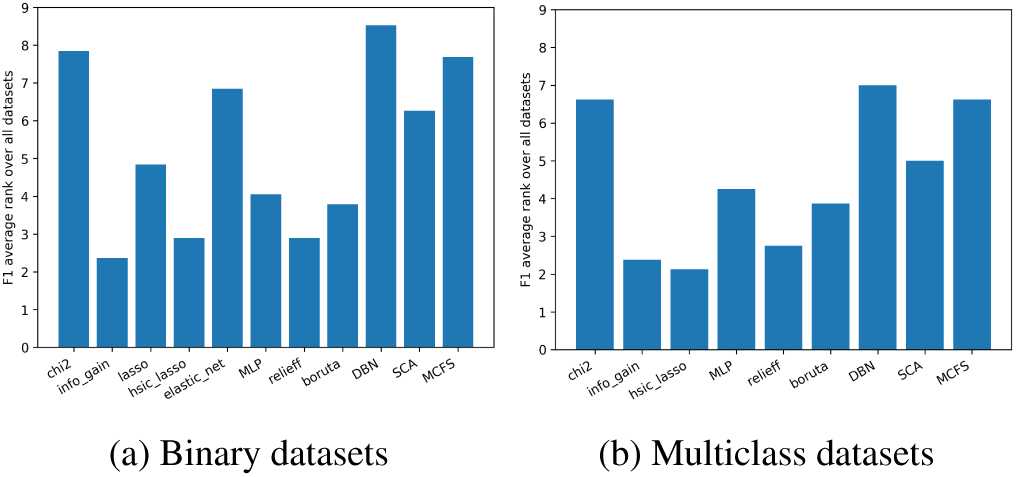
Average mean F1 rank for 11 FS methods classification performance over 19 binary and 8 multi-class classification datasets. The smaller the value, the better.

**Figure 2:**
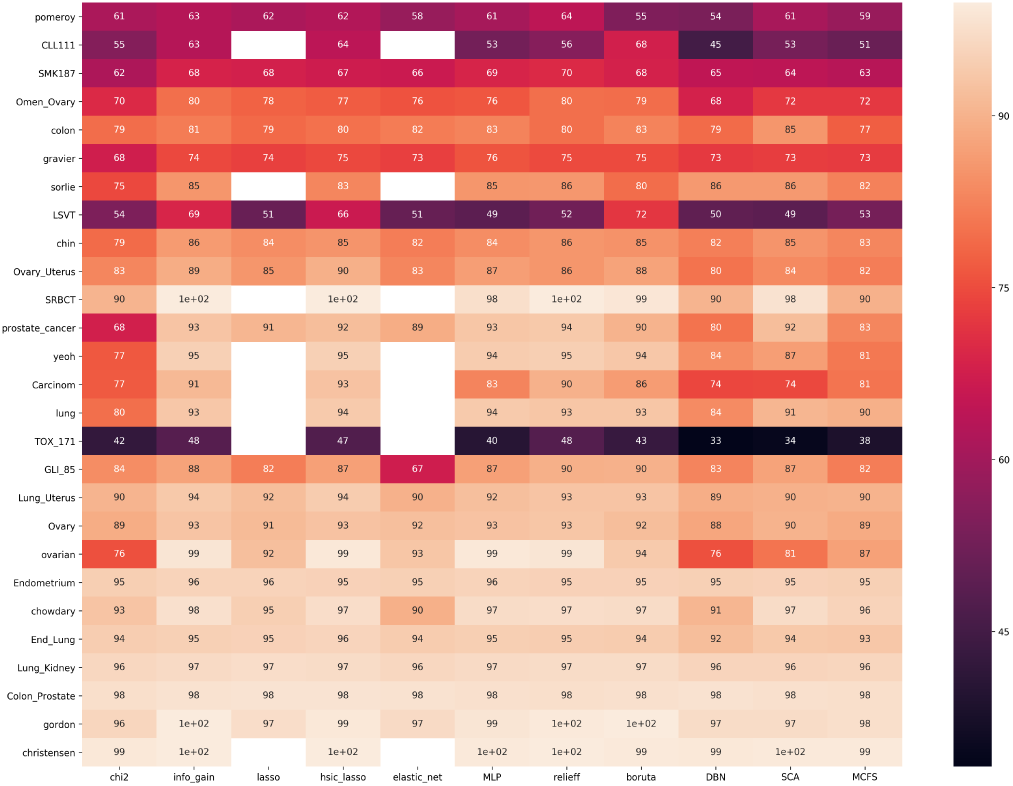
Heat plots for average classification performance of FS methods over all datasets, sorted by N3 complexity measures. christensen has the lowest N3 value while pomeroy has the highest N3 value.

**Figure 3:**
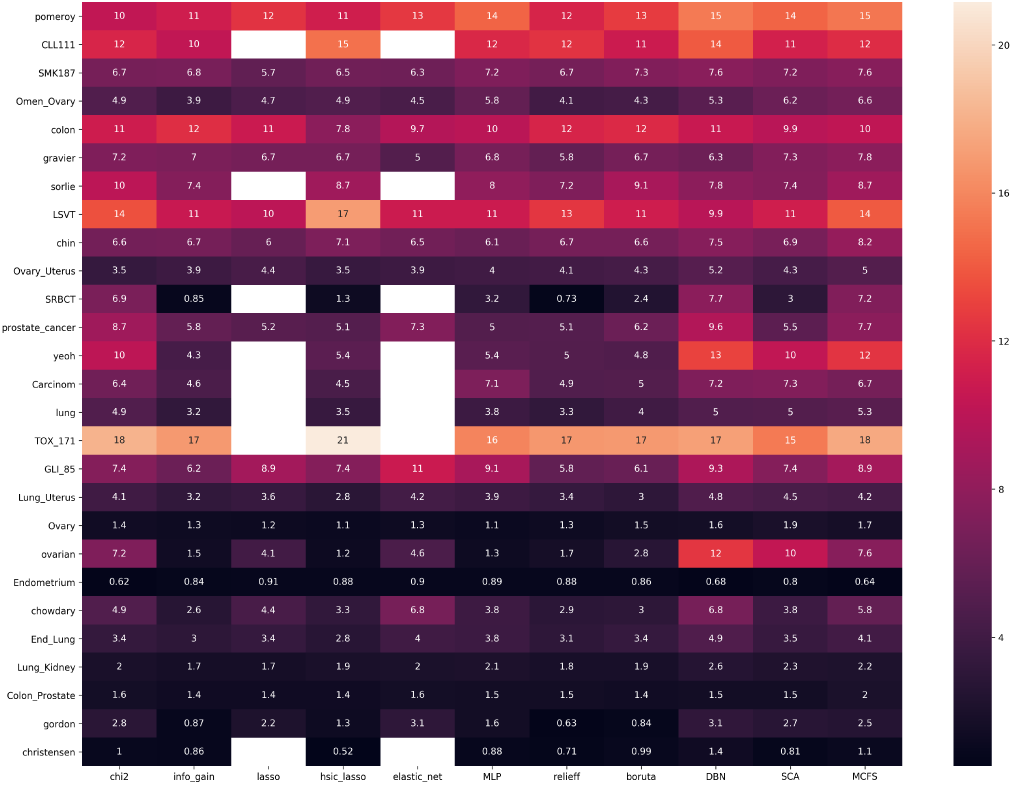
Heat plots for average standard deviation in classification performance of FS methods over all datasets, sorted by N3 complexity measures. christensen has the lowest N3 value while pomeroy has the highest N3 value.

**Figure 4:**
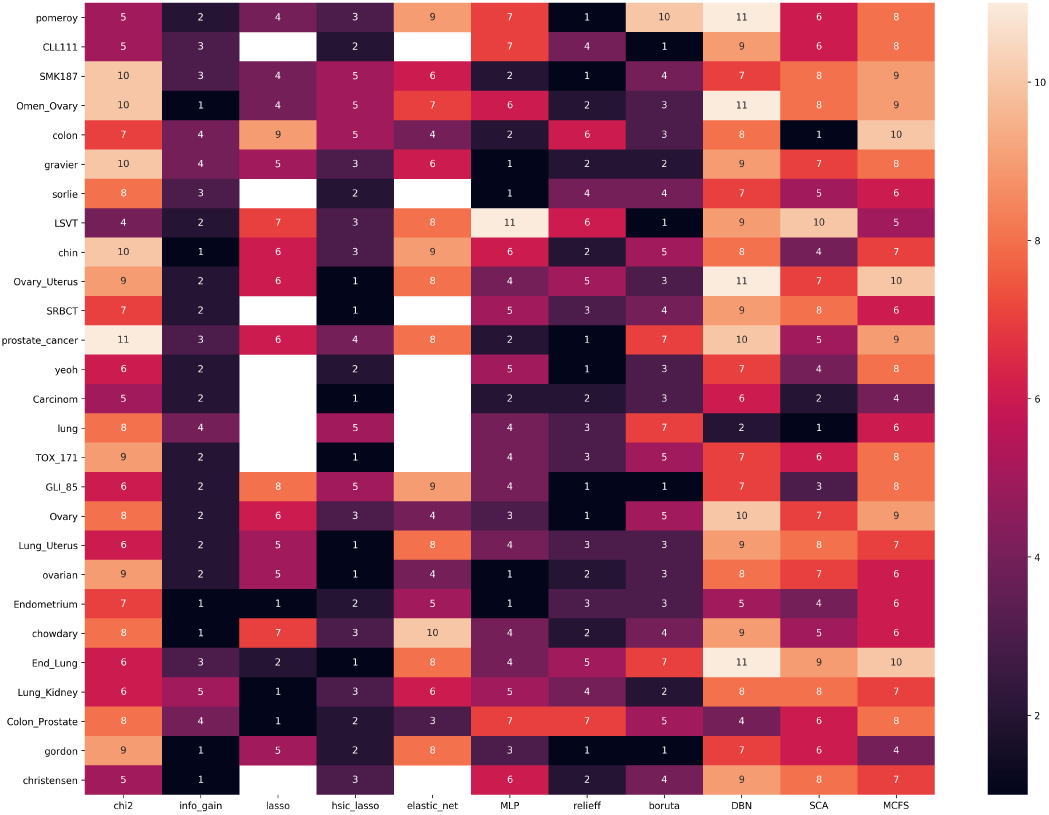
Heat plots for average classification performance ranking of FS methods over all datasets sorted by N3 complexity measures. The lower the rank, the better.

Figure 2 and 4 present the average F1 values and ranking of all feature selection methods over all datasets on a range of number of features (on 30 runs for each number of selected features), respectively.

Looking closely at the plots we see that the average ranking for Info gain over all dataset is smaller than 3. That is to say: most of the time Info gain is in the top 3 performing methods. It is quite surprising that one of the simplest univariate methods tops the list.

Hsic-Lasso is the best-performing method in terms of multiclass classification problem (with 5/8 in the leading position, 2/8 in the second position). In addition, for datasets with small N3 values, Hsic-Lasso is also a good option.

ReliefF is a top-performing method with 6/27 times in the leading position. Despite that, for other datasets, it is not always in the top 3 best performing methods. It is also one of the better performing methods for larger *N* 3 values as compared to the other methods. Note that in case of ReliefF, features are scored based on whether similar feature values are observed in neighboring pairs with the same class labels. High N3 implies that neighboring samples have different classes and the similar features for such pairs would be scored lower. Quantitatively, it shows a lower absolute correlation with N3 as compared to Hsic-Lasso and Info gain.

Even though SCA has the smallest absolute correlation value with N3, it is still on average worse performing than ReliefF even for datasets with smaller N3 values.

Lasso and Elastic Net tend to have similar performance most of the time. However, we observe more variance in Elastic Net.

### 5.3 The returned feature subsets

We ran experiments over a range of number of selected features with the maximum value is 200. However, Boruta and Hsic-Lasso sometimes returned much less number of features even after parameter adjustments. Given the fact that Hsic-Lasso is one of the best performing method while returning a small subset of relevant features, we believe that Hsic-Lasso should be given more preference in the choice of methods, especially in biomarker discovery applications.

We also take a closer look at the set of features returned by different FS methods. At each run, for the number of selected feature equal to 200, we calculate the overlapping portion of the feature subsets returned by different methods. Our hypothesis was that the top-performing method would overlap more and the more similar the method, the larger the overlapping portion of their feature subsets. Though, Hsic-Lasso returns the least number of selected features, around 29% of these returned features overlap with the feature subset returned by other top performing method, ReliefF. Info gain overlap with Boruta (around 54%) and ReliefF (around 47%).

We believe that the questions of whether the overlapping subsets of different feature selection methods enclose the most informative features or not as well as which combination of FS methods might be beneficial to the bio-marker discovery applications are interesting research questions that we will follow in our future work.

### 5.4 Optimal number of selected features

We look at the performance of different FS methods over a range of number of selected features. We observe that FS methods performance does not follow any monotonic trend with regards to the number of selected features. They tend to fluctuate from time to time. There is no fix point where every FS method reach their highest performance. Figure 5 present an example of the FS methods performance on the GLI 85 dataset.

**Figure 5:**
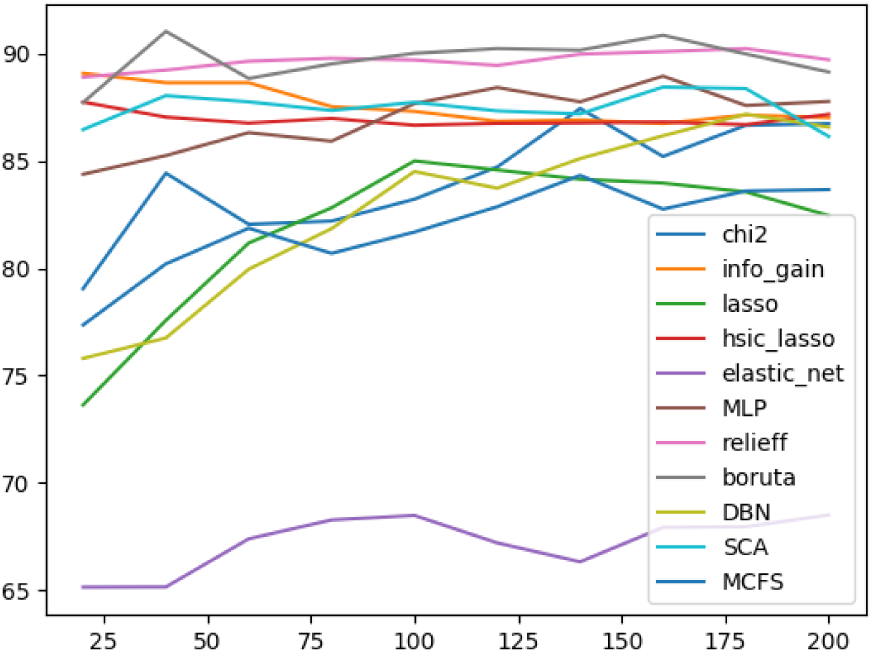
Performance of FS methods over a range of number of selected features for GLI 85 dataset.

We take one step further: for each FS method on a particular dataset, we find the number of selected features at which it attains the highest performance. The result turns out that there is no observed correlation between the optimal number of selected features for FS methods and the datasets’ characteristics. Our hypothesis was built the value of complexity measure *T* 4, i.e., which gives an estimate of the number of features returned by performing dimensionality reduction using PCA (while maintaining 95 % data variability). We believed that all methods would obtain their highest performance by choosing the number of features close to estimate given by T4. Not only this hypothesis turned out to be incorrect, but also the fact that there exists no monotonic trend between the selected number of features and performance, the problem of determining the optimal number of features becomes very hard.

### 5.5 Deep Feature Selection

Even though deep learning methods are usually not recommended for small sample size problems, deep feature selection (DFS) model using MLP shows a relatively good performance (see Figure 1a) and demand further investigations.

### 5.6 Unsupervised Method

Though it is unfair to compare unsupervised method with supervised methods, we included MCFS in our study as as it provided promising performance as compared to other unsupervised methods and sometimes also supervised methods (for example in our initial experiments, we considered a recent unsupervised deep learning method Concrete Autoencoders [1]). We believe that it is a promising method for datasets where the class information might be scarce or not available.

### 5.7 Summary and Recommendations

In the following we summarize our findings and provide recommendations for using feature selection methods.

- For datasets with low Error Rate of Nearest Neighbor classifier(N3), supervised methods Info gain and Hsic-Lasso are recommended to build predictive classification models.
- For data with both low and high N3, ReliefF appears a competitive method.
- Hsic-Lasso and Boruta might return a very small number of relevant features. When we are concerned about both the classification performance and the small number of selected features, Hsic-Lasso should be the best option.
- Different FS methods perform differently with regard to the number of selected features. The points different FS methods reach their highest performance for each dataset varies arbitrarily and neither follow any pattern nor correlate with any of our proposed dataset characteristics.
- In terms of performance, deep learning based methods in general have higher variance in performance than the non-deep learning counter-parts possibly due to smaller sample sizes. Deep Feature selection methods based on MLP, on the other hand shows promising performance and also relatively lower variance.
- The dataset TOX-171 falls out of the normal trend followed by other datasets and despite showing a relatively lower N3 error, it appears to be difficult for all methods.

## 6 Conclusion

In this work we investigated data complexity to understand the suitability of a particular FS method. We conducted an extensive comparative study of 11 FS methods for 27 biological datasets with varying properties. As the optimal number of features is not known in prior, we tested over a wider range where the number of selected features were varied from 20 to 200 with a step of 20. For each number of selected features, we evaluate each method performance 30 times to get a reliable estimate of the method performance. We calculate the correlation between the FS methods performance and the presented data complexity measures. Experimental results show that FS method performance on classification is highly correlated with N3 - a data complexity measures based on the data local neighborhood. We compare 11 FS methods performance in term of average performance, variance and ranking. Summarizing our findings, we also build a recommendation list of various methods. In future we would like to investigate in several directions including a thorough analysis of deep learning models for feature selection, the dependency of optimal number of relevant features on dataset properties and its interplay with method properties and understanding the unusual trend of datasets like TOX-171.

## Footnotes

2 https://CRAN.R-project.org/package=ECoL

3 https://docs.scipy.org/doc/scipy/reference/generated/scipy.stats.zscore.html

4 We use MLP classifier implementation from https://scikit-learn.org/stable/modules/generated/sklearn.neural_network.MLPClassifier.html

5 https://en.wikipedia.org/wiki/Pearson_correlation_coefficient

6 https://docs.scipy.org/doc/scipy/reference/generated/scipy.stats.pearsonr.html

